# Investigating climate-phenology relationships among the most common Italian forest species using Sentinel-2-derived vegetation phenology and productivity products

**DOI:** 10.64898/2026.02.23.707431

**Authors:** Elia Vangi, Giovanni D’Amico, Vincenzo Saponaro, Mattia Niccoli, Gioele Tiberi, Saverio Francini, Costanza Borghi, Alessio Collalti, Francesco Parisi, Gherardo Chirici

**Affiliations:** geoLAB - Laboratory of Forest Geomatics, Dept. of Agriculture, Food, Environment and Forestry, Università degli Studi di Firenze, Via San Bonaventura 13, 50145 Firenze, Italy; Forest Modelling Laboratory, Institute for Agriculture and Forestry Systems in the Mediterranean, National Research Council of Italy (CNR-ISAFOM), Via Madonna Alta 128, 06128, Perugia, Italy; Faculty of Agricultural, Environmental and Food Sciences, Free University of Bozen-Bolzano, Piazza Università/Universitätsplatz, Bolzano 39100, Italy; Department of Architecture (DIDA), University of Florence, Via della Mattonaia 8, 50121 Florence, Italy; Department of Science and Technology of Agriculture and Environment (DISTAL), University of Bologna, 40126 Bologna, Italy; Department of Biosciences and Territory, University of Molise, Contrada Fonte Lappone, 86090 Pesche (Is), Italy; NBFC, National Biodiversity Future Center, Palermo 90133, Italy; Fondazione PerIl Futuro Delle Citta, Florence, Italy

**Keywords:** National Forest Inventory, phenology, remote sensing, SHAP, Copernicus, Machine learning

## Abstract

Climate change is profoundly altering forest phenology and productivity across Europe, with particularly strong impacts in Mediterranean regions characterized by high climatic heterogeneity. Understanding how climatic and site-specific drivers regulate the start, end, and length of the growing season, and how these phenological shifts translate into productivity responses, remains a key challenge for predicting forest carbon dynamics. In this study, we investigate phenological timing and total seasonal productivity across multiple Italian forest species spanning Mediterranean, temperate, and mountain environments, leveraging the new High-Resolution Vegetation Productivity and Phenology product from the Copernicus Land Monitoring Service, machine learning (random forests) modeling, and explainable artificial intelligence analysis (SHAP). Our results confirm a general lengthening of the growing season driven mainly by chilling accumulation and spring temperatures. Warmer conditions advance the start of the season by 1–10 days across species, while the combined effects of temperature, radiation, and moisture can extend the growing season by up to 20–30 days. End-of-season dynamics and season length are more strongly controlled by light and water availability than by temperature alone. In several Mediterranean species, the end of the season can advance by up to 40 days due to summer drought, high vapor pressure deficit, and site exposure. Mediterranean species often show compensatory shifts between season onset and senescence, maintaining a relatively stable length of the season, whereas mountain species exhibit a tighter coupling between delayed onset and shortened season length. Phenological shifts are frequently decoupled from productivity, which is mainly regulated by energy and water availability, highlighting species- and site-specific responses to climate change. The findings of this study highlight the substantial advantage of remote sensing data, coupled with machine learning approaches, for advancing the understanding of forest phenology and productivity across broad spatial and climatic gradients.

## 1. Introduction

Climate change is the leading cause of the global temperature rise since the pre-industrial period. It has already caused a global temperature increase of 0.8 to 1.3°C since 1850 and is expected to rise further to 1.5°C by the middle of the century (IPCC, 2022). For this reason, international negotiations, such as the Paris Agreement and the Glasgow Climate Pact, have set the goal of limiting the rise in temperature well below 2 °C. Yet, turning these duties into actual reductions in atmospheric CO_2_ concentration requires reassessing core carbon accounting principles, integrating them into specific policies and rules, and collecting essential data (Keith et al., 2024). In Europe, climate change impacts are already pronounced and occurring faster than the global average, particularly in Mediterranean and boreal regions (Santini et al., 2014; Noce et al., 2016). The European Union has therefore positioned climate mitigation and adaptation at the core of its policy framework, most notably through the European Green Deal and the legally binding European Climate Law, which commits the EU to climate neutrality by 2050 and to a net reduction of at least 55% in greenhouse gas emissions by 2030 compared to 1990 levels. Achieving these targets requires robust and transparent carbon accounting frameworks, improved monitoring of land-based carbon sinks, and the effective integration of forests and other terrestrial ecosystems into climate strategies. In this context, Europe represents both a critical testbed and a policy driver for translating global climate commitments into operational, data-driven mitigation actions. Among the multitude of key biotic and abiotic climate change indicators, plant phenology is extremely sensitive and reactive to changes in temperature and precipitation patterns (Keenan et al., 2014; Leblans et al., 2017; Kim et al., 2021), making it a perfect fingerprint of climate change impacts (Parmesan, 2003; Peano et al., 2019; Morichetti et al., 2024). Vegetation phenology refers to the timing of recurrent biological events, their causes, and the relationships among them, whether within or across species (Leith, 2013). These events, or phases, include observable periodic changes, such as budburst, flowering, fruiting (including masting), and leaf yellowing and abscission (in broadleaf species), as well as changes that are not directly observable, including the respiration cycle and dormancy in temperate and boreal biomes. Nearly 30 years ago, these phases, which we had long known to be linked to seasonal variability (Piao et al., 2019), were first linked to long-term changes in weather patterns (e.g., Schwartz, 1998; Menzel, 1999). Since then, an increasing number of studies in different biomes and species have acknowledged the effect of climate drift on plant phenology, in particular the earlier spring onset of photosynthesis and the delayed dormancy in autumn, with significant consequences in the carbon and water cycles and energy flows (e.g., Cleland et al., 2007; Garcia-Mozo et al., 2010; Keenan et al., 2014; Diepstraten et al., 2017; Caparros-Santiago et al., 2022; Ren et al., 2024; Morichetti et al., 2024). Phenological changes have repercussions that range from the individual to the community level, modifying the structure of ecosystems and the interactions between species, affecting resistance and resilience, and ultimately compromising their ability to respond to environmental stressors (Visser, 2016; Caparros-Santiago et al., 2022; Gao et al., 2022; Vangi et al., 2024). Plant phenology not only influences natural ecosystems and their inhabitants but also society and economy, affecting all processes directly or indirectly linked to phenological processes, such as timber production, agricultural food supply, and pollen seasonality (Delpierre et al., 2016; Khwarahm et al., 2017). For these reasons, measuring and monitoring phenological phases is crucial to better understand how natural ecosystems respond to increasing climate uncertainty.

Traditionally, monitoring vegetation phenology has relied on field-based observations, often labor-intensive, spatially limited, and time-consuming (Aono and Kazuki, 2008). However, the advancement of remote sensing (RS) technologies has significantly enhanced land surface phenology (LSP) monitoring, bringing it to a more mature and efficient stage, thanks to their high spatial and temporal resolution (Delpierre et al., 2016; Piao et al., 2019; Caparros-Santiago et al., 2022). LSP is usually derived from combinations of spectral values, i.e., vegetation indices (VI), often within the visible and near-infrared (NIR) region, such as normalized vegetation index (NDVI, e.g., Tucker, 1979), enhanced vegetation index (EVI, e.g., Li et al., 2010), photochemical reflectance index (PRI, e.g., Gammon et al., 1992), plant phenology index, derived from both multi- and hyperspectral sensors (PPI, e.g., Jin & Eklundh, 2014) and, more recently, harmonic predictors which help to summarize information related to LSP (Francini et al., 2024).

A structured approach has been established to derive phenological metrics (hereafter referred to as ‘phenometrics’) from RS products. This process involves creating a VI time series (usually NDVI or EVI indices), fitting a function over time (e.g., smoothing functions such as double logistic, Gaussian, or polynomial functions, or Fourier transforms), and extracting relevant phenometrics (Piao et al., 2019; Caparros-Santiago et al., 2022). Some of the most commonly used phenometrics are the day of the year (DoY), which mark the start and end of the growing season (SOS and EOS, respectively). These days mark the start of the green-up phase (i.e., budburst), when greenness begins, cambial activity starts, and the greening-up period ends, corresponding to dormancy (Rodriguez-Galiano et al., 2015). The start and the end of the growing season also define the length of the season (LOS = EOS-SOS). Other typical phenometrics can be derived from the fitted function, from which the minimum and maximum value of VI can be extracted, as well as the amplitude, the slope of the greening and browning phase, and the area under the fitted curve (usually associated with the gross primary productivity [GPP]). Among the multispectral missions, one of the most suitable for retrieving dense and long-term time series of Earth Observation data is the MODIS mission (Zhang et al., 2003; Ahl et al., 2006; Li et al., 2010; see the review by Shin et al., 2023), which offers a temporal resolution of 1-2 days. However, the MODIS sensor has low spatial resolution (ranging from 250 m to 1 km, depending on the bands), making it less suitable for canopy- and species-level applications. Other often-used satellites for monitoring LSP include the Landsat (Melaas et al., 2016) and Sentinel-2 constellations (see the review by Misra et al., 2020), which offer a better trade-off between spatial and temporal resolution across a wide range of applications and spatial scales.

Recently, the Sentinel-2A and 2B time series have been leveraged to produce high-resolution (10 m x 10 m) and high-frequency (five-day) time series of the PPI index at the European level. The PPI is a physically based VI, derived from the radiative transfer equations, and is linearly correlated with Leaf Area Index (LAI, m^2^ m^−2^), having the same measurement unit. It has been shown that LAI is linked to canopy reflectance via the modified Beer’s law, which also holds for the difference vegetation index (DVI = red–NIR) (Jin & Eklundh, 2014). The PPI can be derived from the DVI version of the modified Beer’s law:

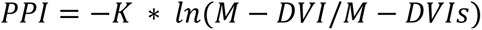

Where DVI is the difference between red and NIR, M is the maximum DVI for a site-specific canopy, DVI_s_ is the DVI of the soil, and K is a gain factor. For a detailed explanation of the PPI formulation and validation, refer to Jin & Eklundh (2014), particularly equations 2-6. The authors have also demonstrated that PPI performs better than NDVI and EVI, particularly in boreal biomes, due to its insensitivity to snow cover and to noise during transition phases.

Thanks to its high linear correlation with LAI, whose changes are among the easiest to observe in the canopy during the growing season, PPI is particularly suited to characterize and monitor phenological phases and, in combination with the spatial scale and frequency of Sentinel-2 observations, make it possible to derive essential phenometrics highly related to physical vegetation variables (such as LAI and GPP). Since 2023, the Copernicus Land Monitoring Service (CLMS) has produced a suite of High-Resolution Vegetation Productivity and Phenology (HRVPP) products, specifically designed to monitor the status and changes in ecosystems, habitats, and land cover (Smets et al., 2024) at local, continental, and global scales. Due to the strong relation with climate factors and biotic and abiotic disturbances, HRVPP products would have great potential to assess future impacts of human activity and climate change on ecosystems, as well as to contribute to reporting land use and land use change (LULUCF) in the framework of international negotiations, ultimately supporting environmental planning from municipality to global level. However, the interactions among phenology, climate, and environmental factors are extremely complex and have been studied in numerous ways. Recent approaches leverage machine learning (ML) algorithms that can capture complex, nonlinear relationships, such as random forests (RF), extreme gradient boosting (XGB), support vector machines (SVM), and various neural network architectures. Common tasks are crop and forest yield estimation (Demisse et al., 2026), species mapping (Yu et al., 2026; Liu et al., 2026), and quantification of climate change impacts (Campioli et al., 2025; Santo et al., 2026). In particular, it is increasingly important to infer causal relationships to understand and mitigate the impacts of climate on forest and agricultural ecosystems. Recently, explainable artificial intelligence (XAI) methods have been developed and used to access the ML black-box decision-making process and support the interpretability of model results. SHapley Additive exPlanations (SHAP) analysis is particularly suited to such aims and is increasingly used in this field thanks to its model-agnostic framework and its ability to compute variable contributions in the same units as the dependent variable (Stumbelj et al., 2014; Lundberg & Lee, 2017; Yu et al., 2026).

Previous studies have successfully used HRVPP products to characterize urban vegetation phenology (Borgongno-Mondino & Fissore, 2022; Ojasalo et al., 2025), monitoring crop phenology (Prikaziuk et al., 2025), and assessing the impact of topography on forest species (Sang et al., 2024). To the best of our knowledge, this study is the first to combine the CLMS HRVPP products, ML, and XAI to investigate forest phenology and productivity at the national level, across multiple species and a wide range of environmental and climatic conditions. For this reason, this study aimed: 1) to assess the potential of the new VPP parameters in characterizing forest species at the national level; and 2) to explore the influence of environmental and climatic factors on the distribution of VPP at the species level. Accordingly, this study sought to advance understanding of forest phenological patterns by evaluating the capacity of the new VPP parameters to characterize species at the national scale and by examining how environmental and climatic drivers shape their distribution across species.

## 2. Materials and methods

### 2.1 Study area

The study focused on the Italian peninsula, covering a total area of more than 300,000 km² and spanning over 11° of latitude, making the area extremely variable and heterogeneous from both environmental and climatic perspectives. Two main mountain ranges cross the peninsula: from north to south, the Apennine range, and from east to west, the Alps, with peaks reaching up to 4000 m asl. Temperate and Mediterranean climates are the primary climate regimes in Italy, followed by an Alpine climate in the mountainous northern regions. More than 35% of the entire territory is covered by forest, according to the 2015 Italian NFI (INFC, 2021), totaling 11,054,458 hectares. Deciduous species represent the dominant forest category (68% of forest area), composed primarily of oak species (*Q. petraea* (Matt.) Liebl*., Q. pubescens* Willd*., Q. robur* L.*, Q. cerris* L.) and European beech (*Fagus sylvatica L.*). Conifer species dominate the seacoast (*P. pinae L., P. pinaster* Aiton) and northern regions (*Pinus sylvestris* L.*, P. nigra* J.F. Arnold*, Picea abies* (L.) H. Karst*, Larix decidua* L.) and form most of the plantation areas. Italian forests are primarily composed of pure broadleaved forests, with pure conifers and mixed forests accounting for just over 10% of the total, except in the Alpine regions, where pure conifer forests predominate. The main silvicultural systems are coppice and high forest, which account for 87% of the total area. The result of these silvicultural systems is the prevalence of even-aged, single-layered forests.

### 2.2. Data

#### 2.2.1 Overview of HRVPP products

The HRVPP suite comprises three different datasets, consisting of (Camacho et al., 2021):

1. VI time series: generated in near-real time at the pixel level, including LAI, NDVI, Fraction of Absorbed Photosynthetically Active Radiation (FAPAR), and PPI.
2. Seasonal trajectory (ST): generated yearly from the 10-day PPI time series by fitting a smoothing and gap-filling function.
3. Vegetation Phenology Parameters (VPP): generated yearly from the ST product, consisting of 13 different phenometrics describing vegetation dynamics during the growing season. Depending on the type of vegetation, up to two growing seasons can be identified.

Figure 1 reports a schematic representation of the VPP product bundle.

**Figure 1:**
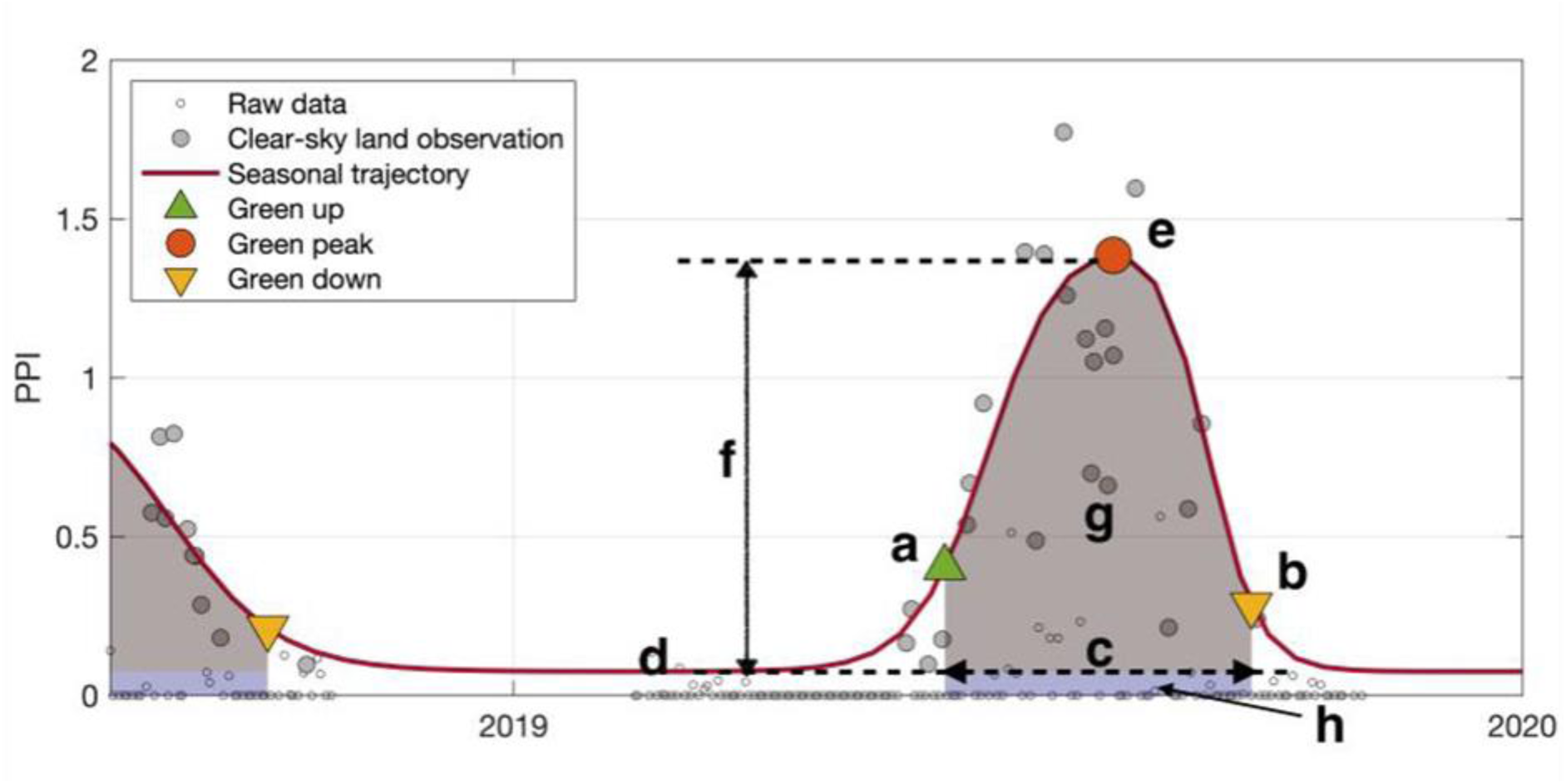
Schematic representation of VPP products (taken from Cai et al., 2023, https://land.copernicus.eu/en/technical-library/algorithm-theoretical-base-document-of-seasonal-trajectories-vpp-parameters/@@download/file)

VPP products are listed in Table 1.

**Table 1:** List of VPP products and their significance with respect to PPI time series.

| VPP |  |  |
| --- | --- | --- |
| Variable name | Letter | Description |
| SOSD | a | Start of the growing season on a specific date |
| EOSD | b | End of the growing season on a specific date |
| LOS | c | Length of the season |
| MAXD | e | Day of maximum productivity |
| MINV | d | Minimum PPI value between two seasons |
| AMPL | f | Amplitude of the season (MAXV–MINV) |
| LSLOPE | a | Slope of the green-up period |
| RSLOPE | b | Slope of the green-down period |
| SPROD | g | Seasonal productivity between SOSD and EOSD minus their base level value |
| TPROD | g+h | Total seasonal productivity between SOSD and EOSD |

All 13 phenometrics are available from 2017 to 2023 (accessed May 27, 2025), and each new year’s product will be provided at the start of the subsequent year (e.g., VPP for 2024 will be available at the beginning of 2025). Below, we provide a brief description of the algorithm and the theoretical basis for these products. For a detailed description of the products and the theoretical basis, refer to Cai et al. (2023).

The HRVPP production process comprises three serial modules. The first module, named BIOPAR-IV, generates VIs (NDVI, FPAR, LAI, and PPI) from time series of Sentinel-2A and -2B images at a 10 m x 10 m spatial resolution. The PPI time series is passed to the next module, TIMESAT, to generate annual seasonal trajectories and VPP products. TIMESAT is a software package for high-resolution time-series analysis (Jönsson and Eklundh, 2004; Eklundh and Jönsson, 2016) that first removes the influence of clouds and shadows from the PPI time series using the QFLAG2 method. After removing clouds and shadows, outliers are identified by fitting a smoothing spline to the raw PPI series and computing the global and local median differences between the smoothed and raw series at each point. An observation is considered an outlier if it exceeds six times the global and/or local median difference. The lower 5^th^ percentile of each pixel in the PPI time series is used as the baseline for the entire series. These values represent the snow-free reflectance of bare ground or stable vegetation. They are used for gap-filling and to assist in the next step, which involves fitting a double logistic function to identify potential seasons that will form the ST products. The double logistic does not assume that the right and left parts of the season are symmetric and ensures that it captures most of the PPI peak, which corresponds to the center of each season. Each potential season is then merged with neighbouring potential seasons until at most two remain. Finally, VPP parameters are derived for each season from the ST. SOSD and ESOD are computed from ST as the DoY in which the PPI reaches 25% and 15% of the ST amplitude, respectively. From SOSD and EOSD, additional parameters are derived, including LOS, SOSV, and EOSV.

#### 2.2.2 NFI data

Field data for this study were collected within the framework of the third Italian NFI, which was formally reported for 2015. The NFI design followed a systematic, unaligned 1 km x 1 km grid sampling with three phases (Fattorini et al., 2006). In the first phase, each sampling point is photointerpreted to assign a land use category. In the second phase, qualitative field data are collected from a subset of points in the forest category. That information includes the silvicultural system, management system, forest category, property, constraints, and accessibility. Finally, in the third phase, a subgroup of the second-phase points, for a total of 6174, is surveyed to acquire quantitative information in 13-m radius circular plots, such as tree species, diameter at breast height (DBH), a sample of tree height, age and current increment, which are aggregated at the plot level and ultimately used to infer different statistics ranging from the NUTS3 to NUTS0 level. In this study, we selected all plots in which the dominant species accounted for at least 80% of the basal area to minimize noise in the phenological response due to mixed forests in the VPP parameters, thereby ensuring a more accurate representation of each species’ phenology. After this initial selection, a total of 91 pure species were identified across 3678 plots. Among these species, we selected those present in at least 100 plots, resulting in 10 species distributed in 2453 plots (Figure 2).

**Figure 2.**
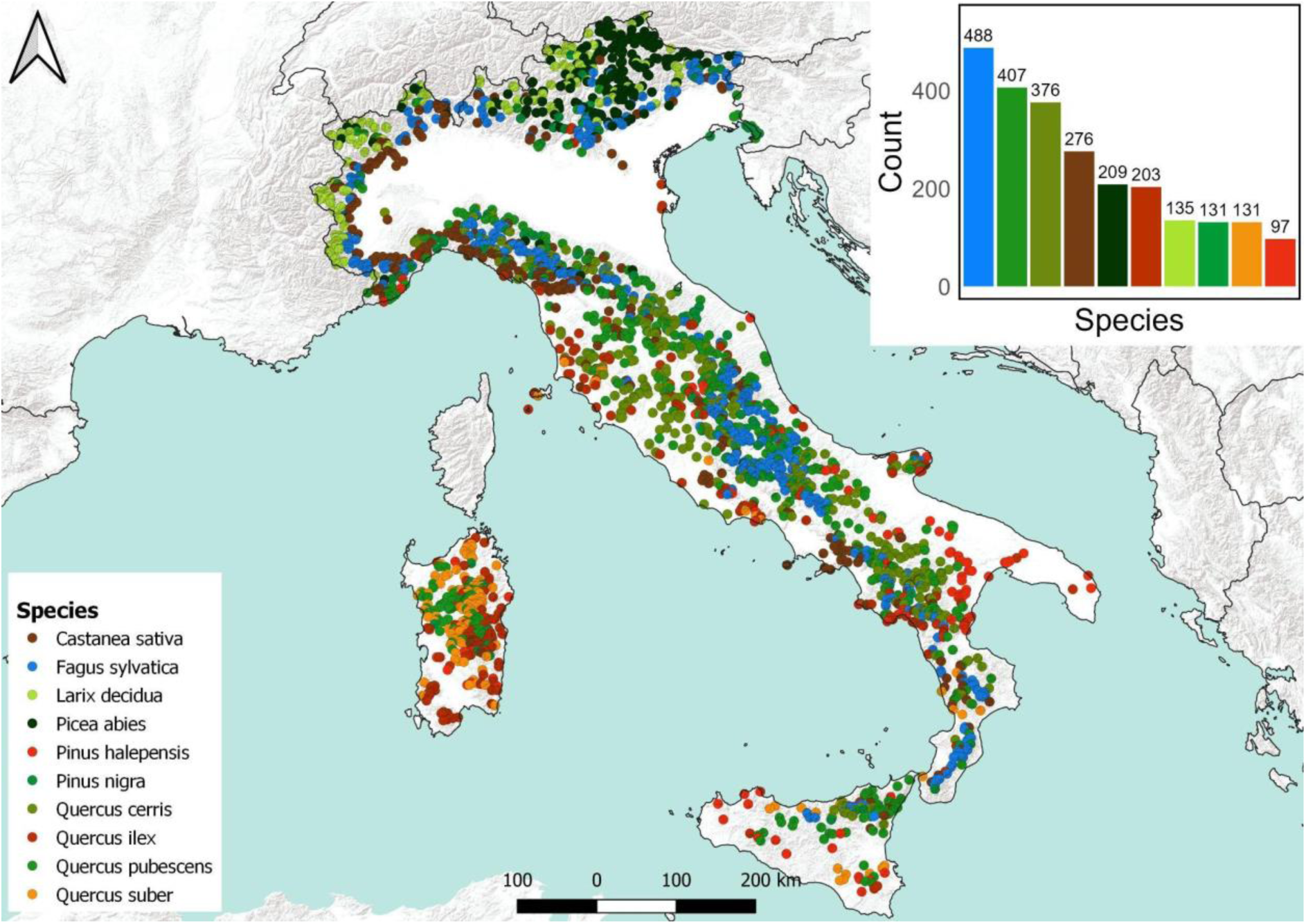
Distribution of the NFI plots analyzed in this study. In the top-right bar plot, the number of plots per species is shown. The bar plot colors correspond to the species indicated in the map legend.

#### 2.2.3. Climate variables

We retrieved climate data from the downscaled ERA5 reanalysis for Italy (Raffa et al., 2021). This reanalysis comprises hourly time series at a higher spatial resolution of 2.2 km for the main climate variables, including air temperature (T), solar radiation (ASOB), precipitation (Pr), vapour pressure deficit (VPD), and dew point temperature (TD), obtained through the Regional Climate Model COSMO5.0_CLM9 and INT2LM 2.06 (Rockel & Geyer, 2008). We downloaded the aforementioned variables for the period 2017-2023 for the entire country. For each year, hourly data were aggregated daily to calculate the minimum, maximum, and mean T (°C), the mean ASOB (MJ m^-2^ d^-1^), the mean VPD (kPa), and the total Pr (mm d^-1^). From hourly T and TD, relative humidity (RH, in %) was calculated (Cai, 2019) and aggregated daily by computing the mean. From the daily climate input, we extracted monthly and seasonal metrics. In particular, the mean, maximum, minimum, range, and sum of each climate variable were calculated for each month and season. Additionally, from the daily average T, we computed the warming rate as the slope of the linear regression of T on time for each month and season. The same procedure was applied for ASOB, and the total Pr. These metrics have been shown to be associated with climate change responses, particularly in the timing of the onset of temperate species (Guralnick et al., 2024). All climate metrics were calculated independently for each year, yielding a total of 176 climate metrics.

#### 2.2.4. Auxiliary variables

Elevation, slope, and aspect were taken from the 10 m resolution digital elevation model (DEM) TINITALY (Tarquini et al., 2007, http://tinitaly.pi.ingv.it/). Additionally, latitude and sea distance were used as auxiliary variables, serving as proxies for precipitation and temperature gradients, which play a crucial role in defining the different phenological phases. These variables represent the local growth conditions for the forest species and remain unchanged throughout their life cycle, unlike climatic metrics. Lastly, we also include stand age as an auxiliary variable, as it is known to affect various physiological processes linked to phenology (Drake et al., 2011; Goulden et al., 2011), including leaf emergence timing, biomass accumulation, and tree species composition. The distribution of these auxiliary variables for each species is shown in Figure S1 (appendix).

### 2.3. Methods

#### 2.3.1. Metric extraction

All 13 VPP parameters were downloaded from the WEKEO data catalog (https://wekeo.copernicus.eu/data?view=catalogue, accessed on May 28, 2025) via the Harmonized Data Access (HDA) API using the R software (version 4.2.1). For each parameter, we downloaded the entire available time series from 2017 to 2023, covering the entire national territory, yielding a total of 128 tiles per parameter per year. The final database comprises 128 tiles × 13 parameters × 7 years, totaling 11,648 tiles and 1.14 TB of data. We selected only the first season for each year and parameter, since seasons are stored chronologically, and we assume that the growing season begins at the start of the year for all Italian forest species. We selected only the seasons that start and end in the same year to ensure comparability among years and species. Each tile was cropped and masked with the Italian national boundary, then reprojected to the common reference system WGS 84 UTM 32N (EPSG: 32632). Five of 13 products were analyzed in this study, specifically related to phenological phases and productivity, namely: SOSD, EOSD, MAXD, LOS, and TPROD. Within a 13 m buffer around each plot location, the mean values of all variables were extracted, including all VPP parameters, climate metrics, and auxiliary variables for each year. For the VPP parameters measured in day of the year (DoY), the mode was extracted instead of the mean. The resulting database consisted of 2453 plots × 7 years = 17,171 observations, including 13 VPPs, 176 climate variables, and five auxiliary variables, for a total of 194 variables. Figure 3 shows the distribution of SOSD, EOSD, and MAXD values across all plots, years, and species.

**Figure 3.**
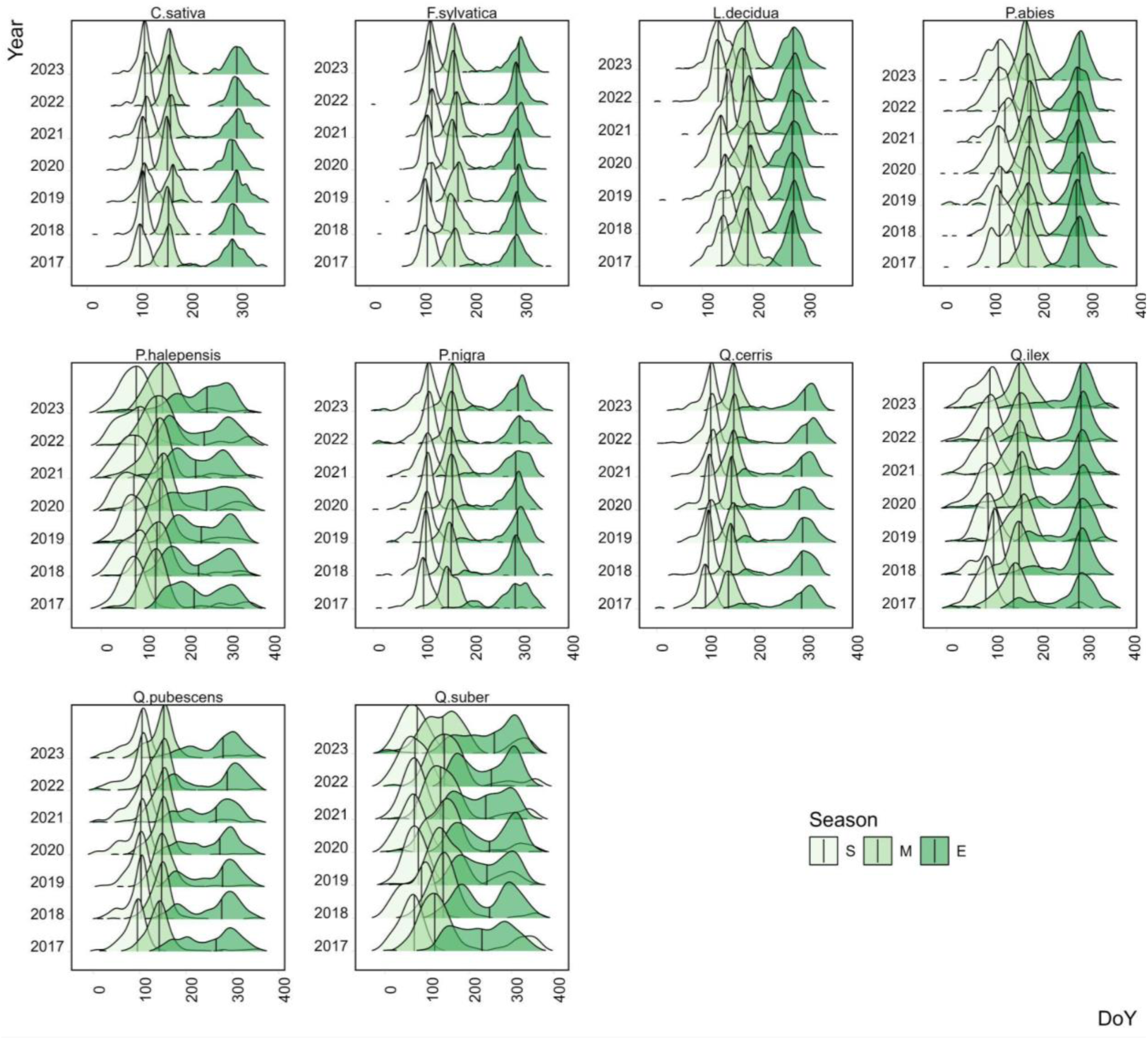
Distribution of phenological parameters for each species and year. Seasons indicate the DoY of the start of the growing season (S), the DoY when the growth peak is reached (M), and the DoY of the end of the growing season (E)

#### 2.3.2. Environmental and climate variable contribution

To determine the significance and contribution of each metric to the phenological phases of the 10 selected species, we used the variable importance (VarImp) from the random forests (RF) algorithm. The original RF algorithm, developed by Breiman (2001), has been shown to be robust to variable autocorrelation and variable scales. It does not assume a linear relationship between the outcome and the predictors, and, thanks to its structure, it can be used to compute a measure of each predictor’s importance. In regression problems, the most common and robust index for importance is the permutation importance, calculated for each predictor as the variation in performance (assessed with an error metric, such as the mean absolute error [MAE] or mean squared error [MSE]) after permuting the chosen predictor and keeping all the other variables unpermutated (Breiman, 2001) in the out-of-bag sample ([OOB], i.e. the sample of data not selected in the tree building). With this method, the relationship between the outcome and each predictor is broken in turn, reducing performance when the outcome is associated with the permuted predictor (Strobl et al., 2007). When selecting variables, the RF permutation-based VarImp offers an advantage over univariate screening methods because it accounts not only for the individual effects of each predictor but also for their interactions with other predictors. However, Strobl et al. (2007) demonstrated that its reliability decreases when predictor variables differ in measurement scales or have varying numbers of categories. Furthermore, importance scores tend to be biased toward variables that are correlated with one another. This limitation becomes especially apparent when combining different predictors, including their interactions, or when predictors of the same type exhibit different category counts within a sample. Different studies have shown how the original RF algorithm may be biased in the variable selection during the split of each tree (Breiman et al., 1984; Kim et al., 2001; Hothorn et al., 2006; Strobl et al., 2007), e.g., by selecting categorical variables with many levels, variables with many possible splits, and/or variables with missing values, seriously affecting the interpretation of the variable’s importance. To address this issue, Genuer et al. (2010) developed a two-step, fully data-driven variable selection procedure based on RF algorithms, called “variable selection using random forest” (VSURF). Below is a brief description of their approach:

The rationale for the approach is to identify, among all important variables highly related to the response variable, the minimal subset that ensures acceptable predictive power when included in a RF model.

##### First step

- *n* RF models with 2000 trees each are fit on the data, and the OOB VarImp is calculated.
- For each variable, VarImp is averaged over the *n* models, and its standard deviation is computed.
- The averaged VarImp are then ranked in decreasing order; a CART model is fit with the VarImp standard deviation as the dependent variable and the ranks as the independent variable.
- Then, only the variables with an averaged VI exceeding the CART model’s minimum prediction are retained. Let’s call this number of variables *m*.

##### Second step: interpretation of variables

- Construct *n’* RF models involving the k first variables, for k = 1 to m, and select the variables involved in the model leading to the smallest OOB error. This leads to considering *m’* variables.

##### Second step: prediction variables

- Starting with the ordered variables retained for interpretation, construct an ascending sequence of RF models by sequentially adding variables and testing them. A variable is added only if the decrease in error exceeds a threshold. The variables of the last model are selected.

The abovementioned procedure is implemented in the R package ‘VSURF’ (Genuer et al., 2015).

To compute the VarImp of the environmental and climatic predictors for predicting VPP parameters, we first averaged annual VPP values at the plot level, along with all predictors. This minimizes the effect of year-to-year climate fluctuations on the species’ phenological signature. Then, for each species, we applied the VSURF approach using, in turn, the VPP parameter as the outcome and all climate and auxiliary variables as predictors. The VarImp for each variable was calculated as the average VarImp resulting from the first step of VSURF. We also report the OOB root-mean-squared error (RMSE) of the model fitted with all variables. The advantage of this method is that the cutoff for determining whether a variable is essential is derived directly from the data itself, without requiring any assumptions. Furthermore, VSURF was designed to be efficient and robust, particularly for high-dimensional (n << p) data with many redundant variables, as in this study.

The subset of variables selected in the prediction step of VSURF was used to calculate the Shapley values for each observation using the SHapley Additive ExPlanation (SHAP) approach.

#### 2.3.3 SHapley Additive ExPlanation (SHAP)

Shapley values, originating in cooperative game theory (Shapley, 1953), have become a prominent method for explaining the predictions of black-box machine learning models and quantifying each feature’s marginal contribution to those predictions. This method is particularly valued for its theoretical properties and model-agnostic nature, making it applicable to a wide range of machine learning models (Stumbelj et al., 2014). The Shapley value is a method for distributing the value of individual predictions into feature contributions. That is, the Shapley values explain the difference between the individual prediction and the global average prediction of a particular model:

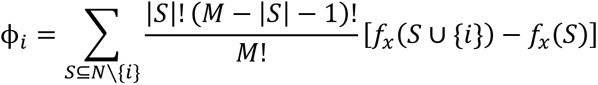

where φ_j_ is the contribution of the j^th^ feature, M is the number of features in the model, S is a subset of features, and v(S) is a contribution function.

In this form, the Shapley value is a weighted average of the differences in the contribution functions across all feature subsets S that exclude feature j (Aas, Jullum, and Løland, 2021).

Lundberg and Lee (2017) have developed the Kernel SHAP method, a practical approach to computing Shapley values in real-world scenarios, since the calculation of (1) may be computationally unfeasible for large datasets, due to the exponential increase in the number of S. Furthermore, the original method was limited to independent features, which is often not the case for remote sensing and climate - derived predictors. Various methods have been proposed to address this assumption (Aas, Jullum, and Løland, 2021; Redelmeier, Jullum, and Aas, 2020; Olsen et al., 2022). In this study, we employed the conditional inference tree (ctree) approach from Redelmeier, Jullum, and Aas (2020), which uses ctree to model each S’s conditional distribution and is implemented in the ‘shapr’ R package (Jullum et al., 2025). SHAP values are expressed in the same unit as the response variable, and their sign and magnitude reveal both the direction and strength of the predictor’s impact on predictions.

A CRF model was fitted to the key predictors identified via the VSURF algorithm, and SHAP values were computed for each prediction for all phenometrics analyzed. Finally, the total contribution of a predictor was calculated as the mean (for the direction) and absolute mean (for magnitude) of its SHAP values.

## 3. Results

### 3.1. Environmental and climate variable ranking

The VSURF underscores that phenology is a multivariate process, modulated by a complex interplay between topography, water availability, and thermal conditions. Generally, climate variables emerged as primary predictors, with temperature and precipitation-related metrics dominating the selection across almost all VPPs. Topographical factors, particularly slope and elevation, were also highly important, being selected by multiple species (e.g., *Castanea sativa, Fagus sylvatica,* and *Larix decidua*) for key periods such as EOSD, LOS, and TPROD, reflecting the strong influence of local microclimate and drainage on vegetation dynamics (Figure 4). This pattern highlights the critical role of early-season growing conditions, in which low temperatures are often the limiting factor for budburst (SOSD), and accumulated winter/spring moisture reserves influence the duration and productivity of the growing season (LOS, TPROD).

**Figure 4.**
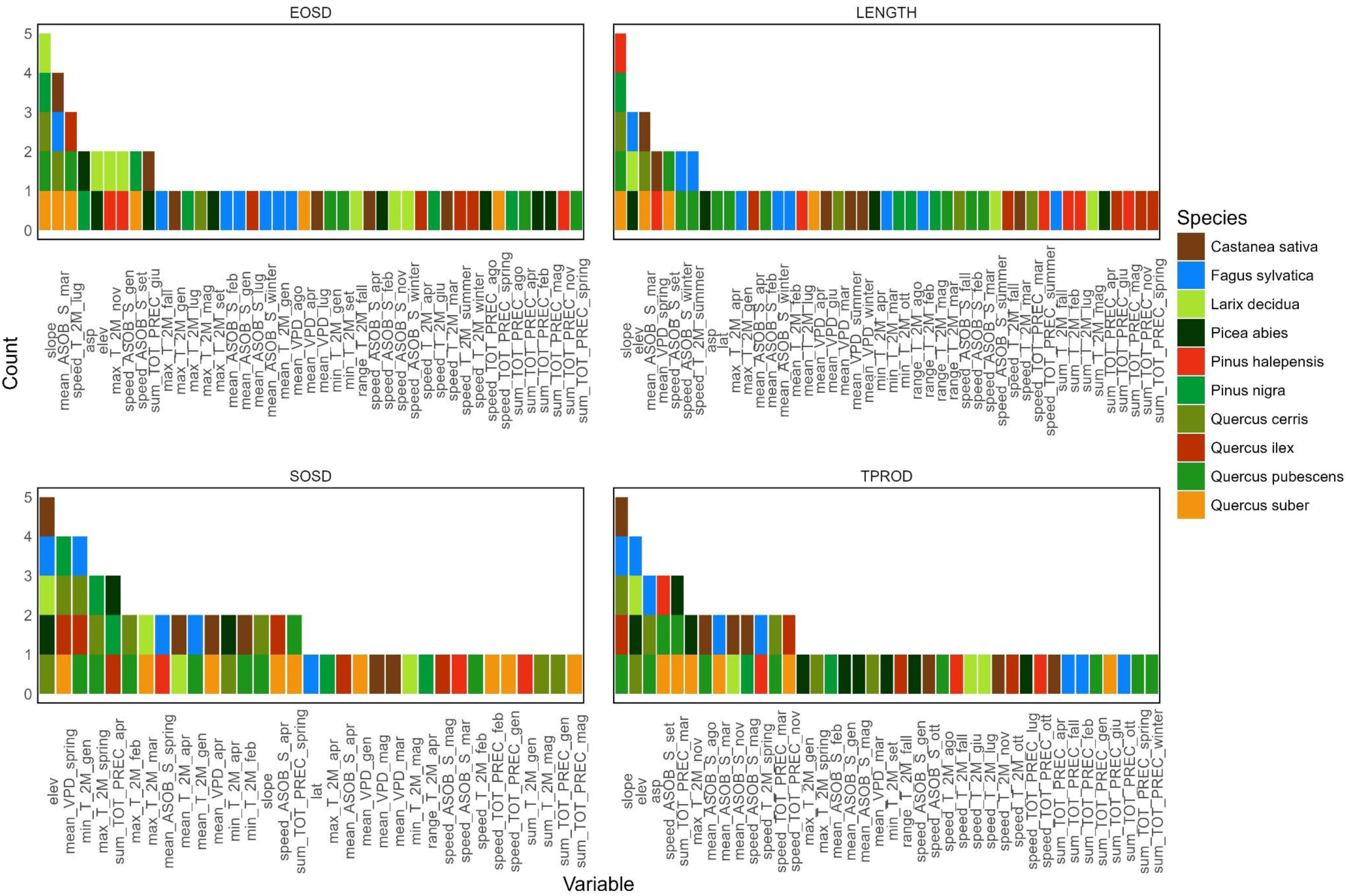
Selected variables in the VSURF procedure for each VPP. Colours map the species for which each variable was selected. EOSD is the DoY of the end of the growing season, LENGTH is the length of the season, SOSD is the DoY of the start of the growing season, TPROD is the total seasonal productivity between SOSD and EOSD.

#### 3.1.1. Start, end, and length of the growing season

The results of the VSURF VarImp in predicting the SOSD for each species are reported in Figure 6. The number of selected variables ranged from three (P. abies) to nine (Q. cerris and Q. suber), with a mean of six across all species. A total of 33 unique variables were selected, with elevation selected in half of the species. Elevation was the variable affecting phenology the most for F. sylvatica, C. sativa, P. abies, Q. cerris, and L. decidua. For other species, the most influential variables were related to winter/spring temperature and radiation (Tmin in January for Q. ilex and Q. pubescens, Tmax in April for P. nigra, the mean spring radiation for P. halepensis, and the total spring precipitation for Q. suber). In particular, T metrics were selected 25 times, followed by VPD and Pr metrics (9 times), and finally ASOB metrics (7 times). The importance magnitude ranged from 1 to 12 days in RMSE. However, it is essential to note that a low magnitude for a single variable does not imply that the variable is unrelated to the outcome; instead, it suggests that its unique contribution to performance is minimal or that it is highly correlated with other variables. This occurs for P. Abies, L.decidua, and F. sylvatica, for which elevation may explain most of the model’s variance. At the same time, different variables are generally highly correlated. In the conditional framework, there is no importance leakage from correlated predictors, so the algorithm will downrank variables that don’t provide unique information beyond what’s already captured. By contrast, most Quercus species exhibited a more uniform decrease in importance across the selected variables, indicating a complex pattern in SOSD that a simple model cannot readily explain. Model’s performance ranges between 8 and 15 days (for F. sylvatica and P. halepensis, respectively), with a mean of 11.5 days among species.

**Figure 5.**
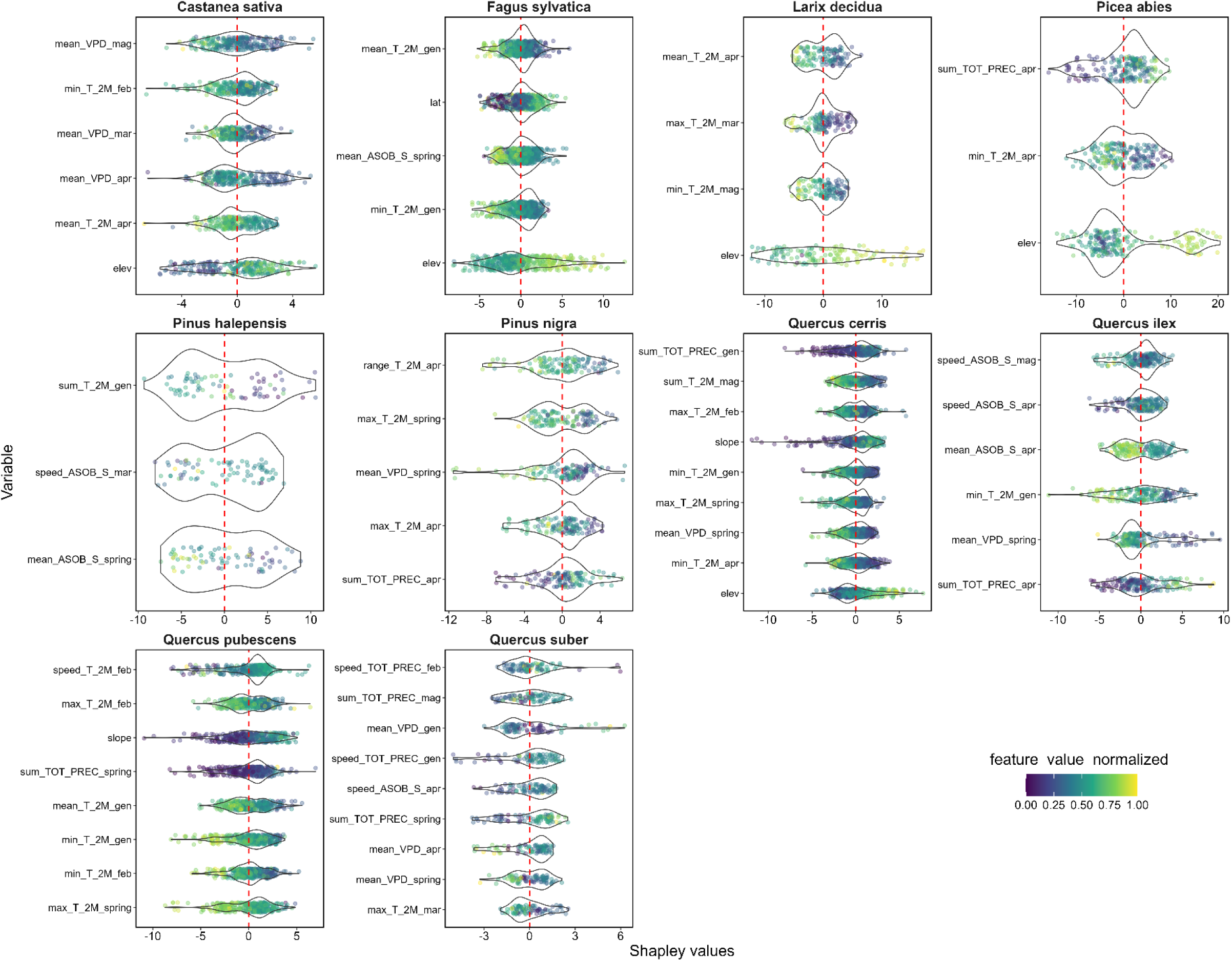
SHAP value (x-axis) distribution of SOSD for the most important variables (y-axis, ordered by increasing importance from top to bottom) selected with the VSURF algorithm for each species. The SHAP unit of measurement is the same as that of the independent variable and represents the difference in days relative to the mean prediction.

**Figure 6.**
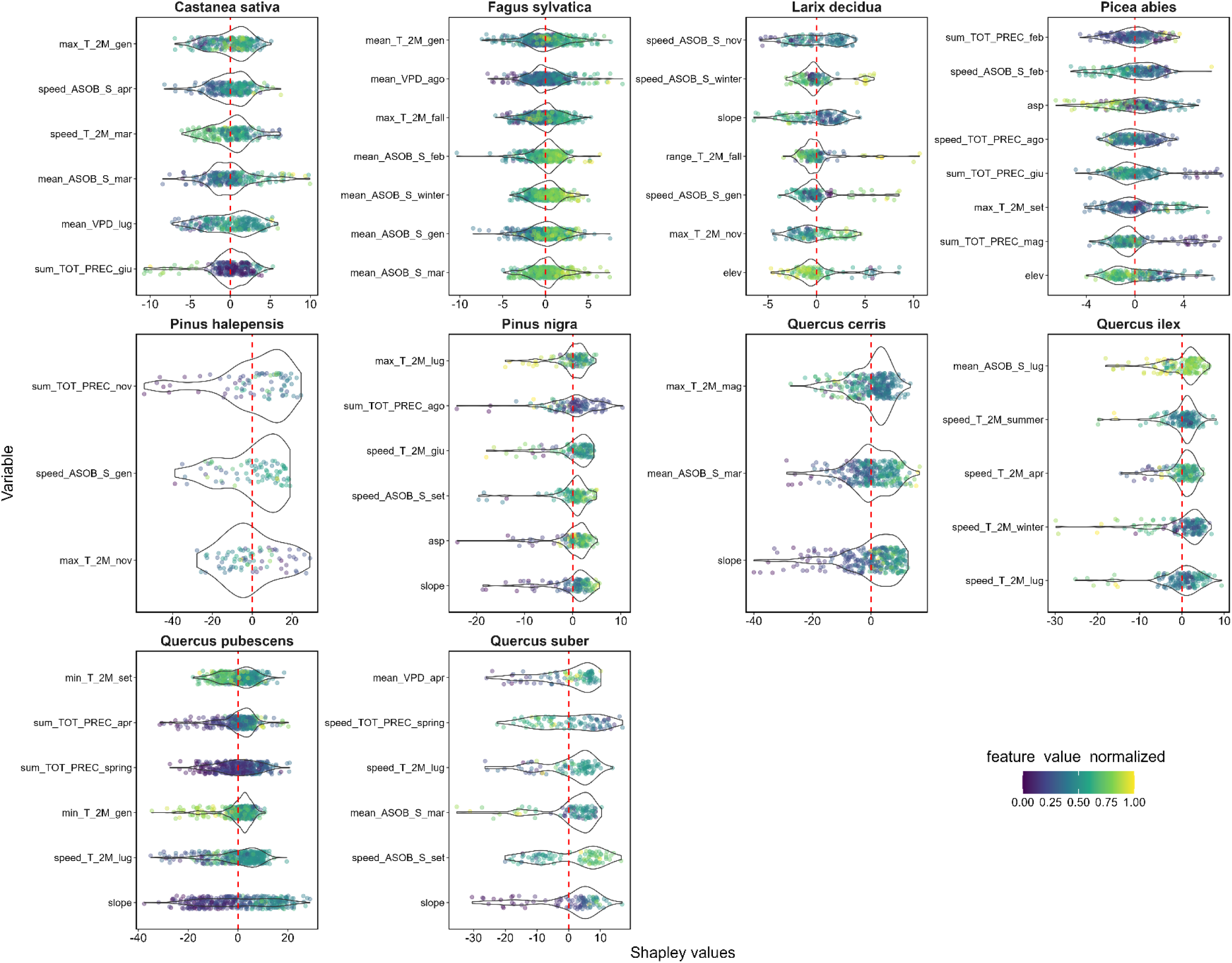
SHAP value (x-axis) distribution of EOSD for the most important variables (y-axis, ordered by increasing importance from top to bottom) selected with the VSURF algorithm for each species. The SHAP unit of measurement is the same as that of the independent variable and represents the difference in days relative to the mean prediction.

Different results were obtained in predicting the EOSD (Figure 7). The number of selected variables ranged from three (P. halepensis) to nine (P. abies), with a mean of six across all species, similar to SOSD. A total of 42 variables were selected. The variables affecting EOSD the most were related to environmental characteristics (the slope for Q. cerris and Q.pubescens, and the elevation for L. decidua), the spring/autumn Pr (P. abies, P. halepensis, Q. suber), and the autumn/winter ASOB (For F. sylvatica and P. nigra). T metrics were selected 19 times, followed by ASOB (16), Pr and VPD metrics (10 and 3 times, respectively), and finally environmental metrics (9 times). The importance magnitude ranged from 1.5 to 14.6 days in RMSE. The model’s performance ranges from 13 to 56 days (for F. sylvatica and P. halepensis, respectively), with a mean of 28 days across species.

**Figure 7.**
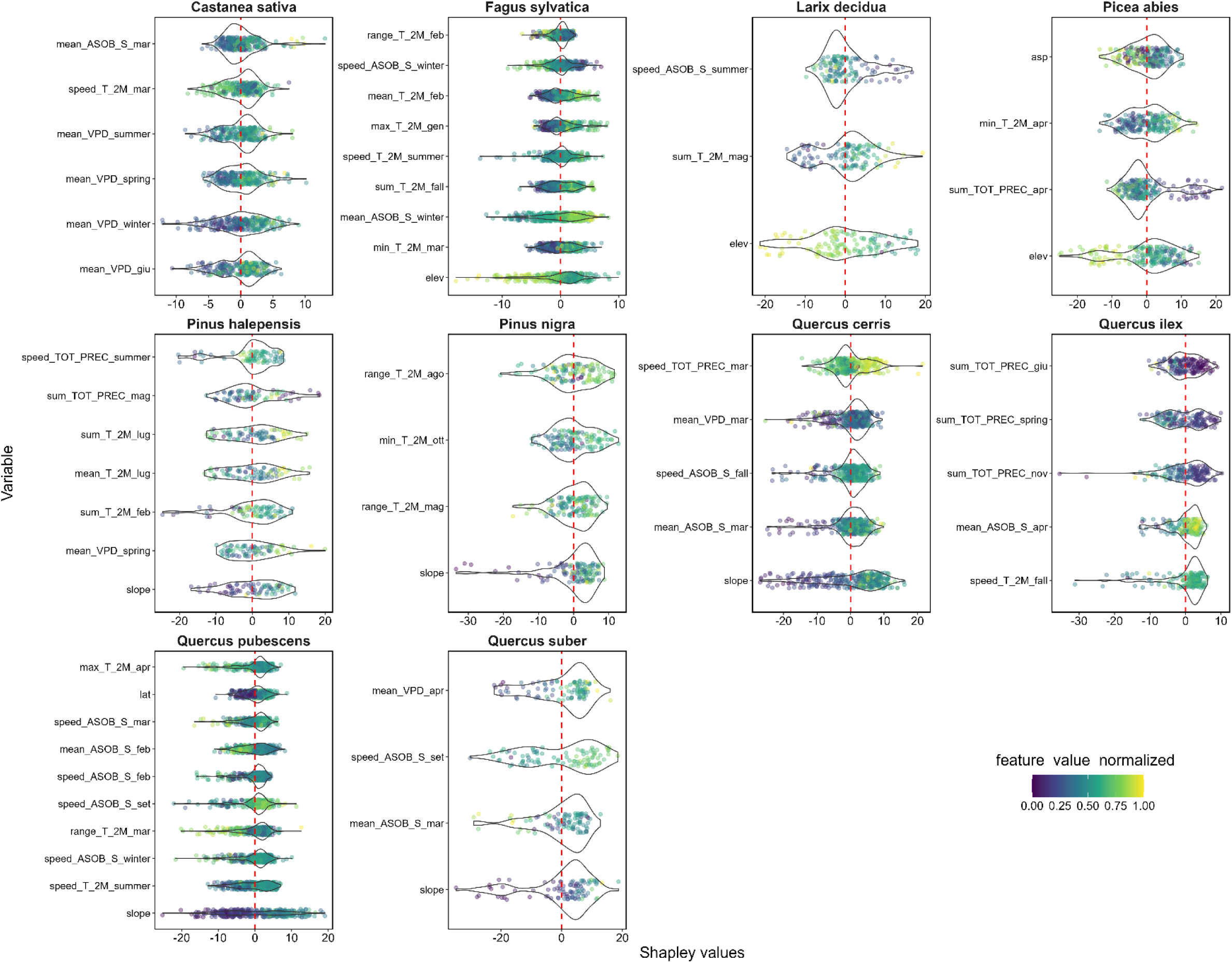
SHAP value (x-axis) distribution of LOS for the most important variables (y-axis, ordered by increasing importance from top to bottom) selected with the VSURF algorithm for each species. The SHAP unit of measurement is the same as that of the independent variable and represents the difference in days relative to the mean prediction.

The most important variables for estimating LOS ranged from 3 (L.decidua) to 10 (Q.pubescens), with a mean of 6 variables. Slope and elevation were the most important variables for six species, followed by autumn/winter T metrics (Q. ilex and P. halepensis) and the speed of ASOB in September (for Q. suber). The magnitude of VarImp ranged between 1.5 and 12 days in RMSE. The model performance ranged from 16 (F. sylvatica, C. sativa) to 50 (P. halepensis) days in RMSE.

#### 3.1.2. Productivity

In the TPROD forecast, the number of selected variables ranges from 2 (P. nigra) to 9 (Q. pubescens and P. abies), with an average of 5.9 across all species. A total of 38 different variables were used, with the slope chosen by half of the species. The most important variables were related to environmental characteristics (the slope for Q. cerris, C. sativa, and Q. pubescens; and the elevation for L. decidua, P. abies, and F. sylvatica), seasonal thermal dynamics, and the spring/autumn Pr (P. abies, P. halepensis, and Q. suber). The model’s performance varies from 46 to 72 in RMSE (for Q. robur and F. sylvatica, respectively), with an average of 58 among species.

### 3.2. SHAP analysis

#### 3.2.1. Start, end, and length of the growing season

The SHAP results for SOSD (Figure 5) indicate that winter and spring temperature variables (minimum and maximum temperatures in January, February, and April) are the primary drivers for most species, confirming the role of temperature in phenological processes. Solar radiation indicators (average intensity and rate of change in April and May) and spring and summer VPD show significant contributions, particularly for Mediterranean species (Q. ilex, Q. suber, P. Halepensis). For mountain species (L. decidua, P. abies, and F. sylvatica), elevation has a positive impact (high elevation favours the start of the season). At the same time, topographic variables such as slope emerge as secondary but sometimes significant factors for some oak species (Q. cerris and Q.pubescence).

In the analysis of EOSD (Figure 6), winter and spring thermal variables (minimum and maximum temperatures in January, February, and April) confirm their role as determinants for most species. Solar radiation indicators, such as spring radiation speed and average, spring and summer vapor pressure deficit (VPD), and precipitation, are particularly influential for Mediterranean species (Q. ilex, Q. suber, P. halepensis). Elevation has a primary impact on mountain species (Larix decidua and Picea abies), with solar radiation indicators of relative importance.

The SHAP analysis reveals that for LOS, the variables are temperature, water availability, and vapor pressure deficit (VPD), which are determining factors for most Mediterranean species (Q. ilex, Q. suber, and P. halepensis). Summer precipitation and spring VPD significantly affect the duration of the season in C. sativa. Mountain species (L. decidua and P. abies) exhibit strong elevational and summer solar radiation dependencies, confirming their preference for cool, bright environments. Slope is a limiting factor for P. nigra, Q. cerris, and Q. pubescens, reducing seasonal length.

#### 3.2.2. Productivity

The SHAP-based models for TPROD show that solar radiation, particularly the average and rate of change in spring and autumn, is a major driver of species such as P. nigra and C. sativa, indicating a strong dependence on energy availability. For Mediterranean species (P. halepensis, Q. ilex, Q. suber), productivity is influenced by seasonal thermal dynamics (rate of temperature change) and the distribution of autumn and spring precipitation, with patterns suggesting thresholds beyond which the effect becomes negative. For mountain species (P. abies, L. decidua, and F. sylvatica), elevation remains a critical factor. Slope remains a limiting factor for several species, particularly Q. cerris and Q. pubescens. Moreover, for species such as F. sylvatica, Q. cerris and Q. pubescens, aspect is one of the key factors that could influence sunlight or microclimate conditions.

**Figure 8.**
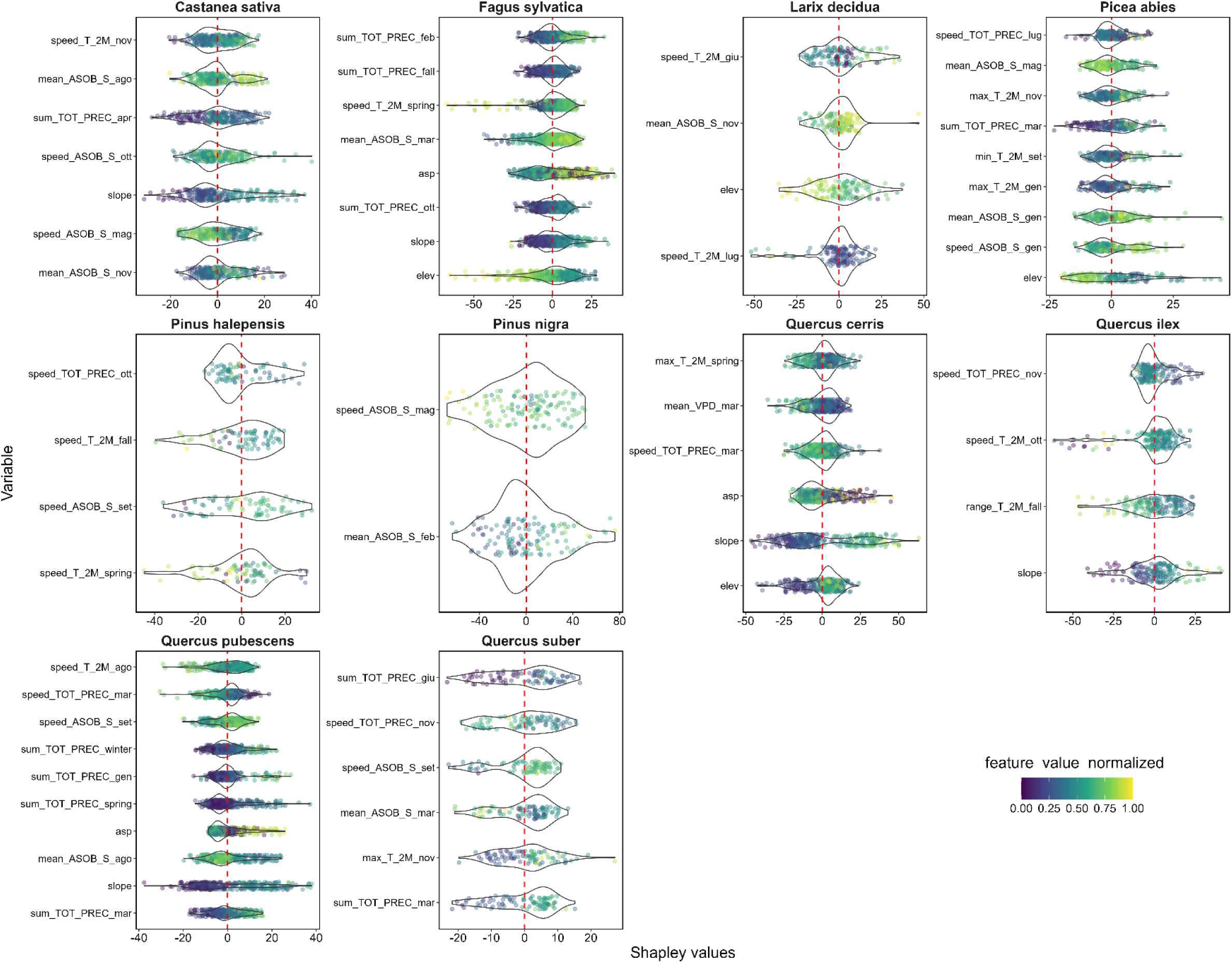
SHAP value (x-axis) distribution of TPROD for the most important variables (y-axis, ordered by increasing importance from top to bottom) selected with the VSURF algorithm for each species. The SHAP unit of measurement is the same as that of the independent variable and represents the difference in PPI^2^ relative to the mean prediction.

## 4. Discussion

### 4.1. Start, end, and length of the season

Climate change is markedly altering weather conditions, especially in Mediterranean areas. Increases in mean air temperature, a higher incidence of severe heat waves, extended periods of drought, and altered precipitation patterns are lengthening the growing season and collectively redefining the regional climate regime, as the carbon and water cycle. These climatic shifts have influenced the phenology not only of Mediterranean species but also of other mountain species, such as European beech (Nezval et al., 2025), Norway spruce (P. abies, K. 1881), and Scots pine (Pinus sylvestris L., 1753) in central and northern Europe (Vangi et al., 2024a; 2024b). Physiological status and phenological phases are primary factors driving forest biomass accumulation and carbon sink capacity, which are critical ecosystem utilities for climate change mitigation. In Italy, due to extreme variation in climatic and environmental variables, individuals of the same species face very different conditions, enabling the assessment of phenological responses to environmental variability.

One observable symptom of the change is an increase in LOS, linked to early budburst and late dormancy (Liu et al., 2018; Jiang et al., 2022; Nezval et al., 2025), driven by rising spring and autumn temperatures. Many studies have observed that leaf unfolding depends on environmental variables, such as air temperature, solar radiation, soil moisture, and elevation (Salehi et al., 2020; Vogel, 2022; Skvareninova et al., 2024). In central Europe, for important broadleaf species such as the European beech, rising temperatures and the CO_2_ fertilization effect are the main drivers of longer growing seasons, leading to increased productivity (Hájková et al., 2010; Skvareninova et al., 2024; Vangi et al., 2024). As said, individuals who grow in Mediterranean areas tend to be more susceptible than those in Alpine and mountain areas.

Our results confirm this common behaviour across many Italian species: individuals exposed to higher winter and spring temperatures (winter chilling) anticipate the SOSD by 1 to 10 days, thereby lengthening the growing season by up to 10 days solely due to temperature. We found that the trend in minimum temperature was more influential in shifting the SOSD than the average and maximum temperatures, confirming the role of chilling temperatures in quiescence and budburst (Hänninen, 2016). The frequent selection of minimum temperatures in winter months (January–February) and maximum temperatures in April suggests that both chilling accumulation and spring forcing are relevant, a pattern widely reported for both broadleaf and conifer species. When other climatic drivers are taken into account, the growing season can be extended by 20-30 days in many considered species (e.g., Q. ilex, Q. pubescens, Q. cerris, P. halepensis, L. decidua, P. abies), reinforcing the findings of Nezval et al. (2020; 2025) in the Czech Republic. Other sites’ characteristics act as proxies for climatic drivers, such as elevation and latitude, thereby shifting the SOSD along climatic gradients, especially in Alpine species, where temperature and snow persistence determine the length of the growing season. This may explain the relatively few individual climatic variables selected for these mountain species in the VSURF framework, which likely reflects high collinearity among temperature, radiation, and moisture variables along elevational gradients. In such cases, elevation effectively captures most of the variance relevant to SOSD and LOS, downweighting other correlated predictors despite their known physiological relevance and widespread species (i.e., Q. cerris and C. sativa, which vegetated along some of the widest altitudinal gradients in Italy). In contrast, Quercus species exhibit a more complex and plastic phenological response, with greater interannual variability, suggesting that more factors jointly regulate the onset of seasons.

The EOSD and LOS seem to be regulated more by light and water availability (both in the form of air and soil humidity and precipitation, and their proxies, such as the site’s slope and aspect), and their effects tend to be greater than those of regulating the SOSD. In fact, EOSD can be anticipated by up to 40 days in certain sites and species (Q. pubescens, Q. cerris, P. halepensis). We observed different growth-regulation strategies among the studied species. Mediterranean species often display compensatory shifts between the start and end of the growing season, resulting in relatively stable season lengths. This behaviour is consistent with previous studies indicating that phenological plasticity in Mediterranean ecosystems is constrained by summer drought and atmospheric water demand (Delpierre et al., 2016; Caparros-Santiago et al., 2022), rather than by spring and autumn temperatures and light availability alone. On the contrary, mountain species such as L. decidua and P. abies (even F. sylvatica at higher latitudes and elevations) exhibit stronger coupling between delayed season onset and reduced season length, reflecting the dominant role of temperature and energy limitations, as also reported by Keenan et al. (2014). However, it is worth noting that there is no consensus on which factors drive EOSD (Campioli et al., 2025; Nezval et al., 2025), as evidenced by the model’s generally lower performance in predicting it. Evidence suggests that trees initiate the cessation process well before observed phenological changes (Smith, 2000). From our data, it appears that summer conditions (in particular, precipitation amounts and the rate of summer heating) already play an important role in determining the leaf senescence (i.e., for C. sativa, P. abies, Q. ilex, and Q. pubescens), usually by retarding the EOSD (but the shift is site and species-dependent), as found by other studies (Dox et al., 2022; Xie et al., 2018). Furthermore, data suggest that spring temperature can also play a role in leaf senescence, as hypothesized by Zohner et al. (2023), reinforcing the idea of yet-unknown intra-seasonal legacy effects.

Other species exhibit a more complex interplay among climatic variables and across different seasons: while temperatures are suggested to control the rate of senescence (Gill et al., 2015; Zohner et al., 2023), and indeed it emerges as a selected predictor for numerous species, the trigger of the onset of senescence is less clear. In our study, solar radiation was selected across all species except one (Q. pubescens), suggesting its role as a possible trigger of leaf senescence. It is interesting to note that in mountain and Alpine species, radiation was selected as the rate of change (a proxy for the photoperiod), which has been hypothesized to be a more important factor at high latitudes and altitudes (Jiang et al., 2022) than radiation intensity, which instead regulates senescence in many European deciduous species (Liu et al., 2023). Also, water availability (soil and air moisture and precipitation) can trigger stomatal closure and shorten the LOS, as observed in P. halepensis, Q. Suber, and Q. ilex (strictly Mediterranean species). Autumn phenology is a tightly controlled process of cellular deconstruction and recycling, making its dynamics intrinsically more difficult to predict than the spring phenology, which appears thermodynamically controlled (in many forest models, the start of the season is determined by the Growing Degree Days, a measure of heat accumulation, e.g., Collalti et al., 2016; Dufrêne et al., 2005; Collalti et al., 2016; Zhou et al., 2023).

### 4.2. Productivity

A common assumption in climate change studies is that phenological shifts, particularly earlier spring onset and longer growing seasons, lead to increased ecosystem productivity and carbon uptake (Keenan et al., 2014; Piao et al., 2019). However, the results of this study reveal a more complex relationship, with phenological timing and productivity often decoupled. Several species show significant interannual differences in the timing of the season’s start or end without corresponding changes in total seasonal productivity, while others exhibit productivity variations in the absence of marked phenological shifts. Similar non-linear and context-dependent relationships have been reported in recent studies (Ren et al., 2024; Morichetti et al., 2024; Andreatta et al., 2025), highlighting the role of limiting factors such as VPD, precipitation seasonality, and light availability. However, productivity depends not only on phenological phases but also on physiological traits, disturbance history, and site conditions, which explain the generally lower predictive performance of the models for TPROD (Saponaro et al., 2025).

As for phenological phases, TPROD seems to be regulated primarily by water and energy availability in Mediterranean species, particularly spring-autumn precipitation and autumn solar radiation intensity, as Liu et al. (2024) have found for about 40% of European broadleaved species. We observed nonlinear relationships between productivity and these variables, with many exhibiting threshold effects. For instance, the rating of the change in solar radiation in autumn has a positive effect only when it is slightly negative (i.e., the slope of the linear regression > -2), the same applied for the heating rate of autumn. Aspect exhibits a similar behaviour, being positively influential from 0° to 90° and from 250° to 0° (indicating that, on the most exposed slopes, temperatures and solar radiation can become limiting factors). Aspect can play a key role in shaping local microclimatic conditions, particularly beneath the forest canopy, by reducing the variability of air and soil temperatures and buffering the impacts of climate change (Rita et al., 2021). On the contrary, precipitation and temperature, regardless of the metric and the period, are always positively correlated with TPROD. Several studies on xylem and phloem phenology agreed that a warming climate would result in a lengthening of the growing season; however, when accounting for drought effects, this extension may be offset by a reduction in the period available for wood formation, cell enlargement, and secondary cell wall thickening (Perez-de-Lis et al., 2017; Dox et al., 2020; Redemacher et al., 2021; Campioli et al., 2025).

Alpine and mountain species once again show a dominant elevational signal, reinforcing the idea that productivity in these ecosystems is constrained by temperature regimes and growing season length rather than by water availability.

It is worth noting that climatic conditions are not the only factors in determining phenological processes and their coupling with physiological responses, such as radial growth and biomass accumulation. As a matter of fact, different species in the same or similar environments show very different behaviour, as found, for example, by Camarero et al. (2010) and Cuny et al. (2012). Different strategies among species, and even different yearly weather, affect tree carbon allocation, favoring or hindering the early development of leaves, productivity, and, therefore, growth, as well as delaying leaf abscission and dormancy (Vicente-Serrano et al., 2016).

## 5. Limitations and future directions

Despite the advantages of remote sensing monitoring at large spatial scales and a machine-learning-based approach, coupled with a recently developed XAI technique, several limitations should be acknowledged. A primary limitation concerns the indirect nature of remotely sensed phenology. Vegetation indices do not directly observe phenological events (e.g., budburst, leaf senescence, or cambial reactivation) but rather capture changes in canopy optical properties that integrate multiple biological and structural processes, including understory phenology. As a result, the detected start or end of the season may lag behind or precede true physiological transitions, particularly for processes occurring below the canopy, such as xylem and phloem activity. To limit this discrepancy, we have selected field plots dominated by a single species (basal area>80%). However, spectral mixing due to spatial resolution, differences in species or age cohorts, snow cover, clouds, and shadows is extremely difficult to control.

Another limitation is the inherently complex, multi-causal nature of autumn phenology (senescence, leaf abscission, lignification), which is influenced by factors not explicitly captured here, such as interannual legacy effects, soil properties, biotic interactions, and disturbance history.

Finally, aggregating VPP over 2017-2023 reduced interannual variability in phenophases at the cost of losing information about specific short-term yearly extreme events, which are increasingly relevant to forest functioning. Extending the analysis to longer time series and explicitly incorporating climate extremes would improve the robustness of predictions under future scenarios.

Future research should integrate mechanistic variables, including soil moisture dynamics, snow-cover duration, and physiological indicators of stress, to better understand end-of-season processes and their drivers. Furthermore, coupled with current-year conditions, previous-year climatic factors and phenophases also affect current-year phenology (Dox et al., 2022; Gu et al., 2022). Intra- and interannual climate variability, as well as their legacy effects, should receive greater attention in future research.

From a broader perspective, integrating phenology-based remote sensing products with the emerging framework of satellite biodiversity represents a promising future direction. Long-term, high-resolution phenological metrics, combined with information on vegetation productivity and structural complexity, can serve as spatially explicit proxies for habitat heterogeneity and biodiversity patterns across taxa. Coupling phenological indicators with biodiversity-relevant EO products (such as phenological variables) could support integrated assessments of climate impacts on ecosystem functioning, carbon cycling, and biodiversity.

Finally, coupling data-driven approaches with process-based models could help resolve discrepancies between observed canopy-level signals and underlying physiological processes, improving predictions of end-of-season dynamics, and linking phenological shifts to productivity and carbon cycling, especially in the context of climate change.

## 6. Conclusion

This study is among the first to apply the Copernicus High Resolution Vegetation Phenology and Productivity (HRVPP) products to investigate forest phenology and productivity across multiple species and bioclimatic contexts. We integrated ML-based variable importance with SHAP analyses to jointly preserve predictive performance and ecological interpretability across multiple species on a wide environmental gradient. This dual framework allows the identification of both dominant drivers and nonlinear responses. Moreover, the inclusion of diverse bioclimatic contexts, from Mediterranean to montane species, enables a comparative assessment of species-specific sensitivities to climatic and topographic constraints.

Phenological models and RS monitoring bridge gaps in observations with limited spatial or temporal coverage by effectively scaling field information, such as transect surveys or phenocams. Since the recognition of climate-driven shifts in phenology, there has been increasing interest in collecting long-term time series, which has enabled modeling of phenological patterns and trends, thereby enhancing our understanding of tree physiology, carbon assimilation, species distributions, and disturbance dynamics (Silvestro et al., 2025). By demonstrating the applicability of HRVPP products for species-level phenological analyses, this work represents a substantial improvement over coarser-resolution datasets traditionally used for land-surface phenology analyses (Misra et al., 2020).

Given the increase in temperatures, the decrease in precipitation and relative humidity in the period 2017-2023 compared to the period 1981-2005 (Figure S2, Appendix), and the results of this study, it can be inferred that Italian forest species are undergoing important physiological changes (changes in phenological phases and productive cycles), trying to adapt to a fast-changing environment. Ecologically, the findings suggest that future warming may continue to advance SOSD across species, but its effects on EOSD, LOS, and productivity will be far more species- and site-specific, particularly in water-limited environments. This complexity has important implications for forecasting forest responses to climate change and for the parameterization of dynamic vegetation and ecosystem models.

## Acknowledgments

E.V. acknowledges NextGenCarbon H2020 project funded by the European Commission, number 101184989 call HORIZON-CL5-2024-D1-01-07 and the Space It Up! project funded by the Italian Space Agency, ASI, and the Ministry of University and Research, MUR, under contract n. 2024-5-E.0 - CUP n. I53D24000060005. D.D. and A.C. acknowledge the project funded under the National Recovery and Resilience Plan (NRRP), Mission 4 Component 2 Investment 1.4 - Call for tender No. 3138 of December 16, 2021, rectified by Decree n. 3175 of December 18, 2021 of Italian Ministry of University and Research funded by the European Union – NextGenerationEU under award Number: Project code CN_00000033, Concession Decree No. 1034 of June 17, 2022 adopted by the Italian Ministry of University and Research, CUP B83C22002930006, Project title “National Biodiversity Future Centre - NBFC and the project FORESTNAVIGATOR Horizon Europe research and innovation program under grant agreement No. 101056875.

## Data availability

The HRVPP and climate data are freely available from their original sources. All other data, images, and codes are available via the Zenodo repository: xxxx

## 7. Appendix

**Figure S1.**
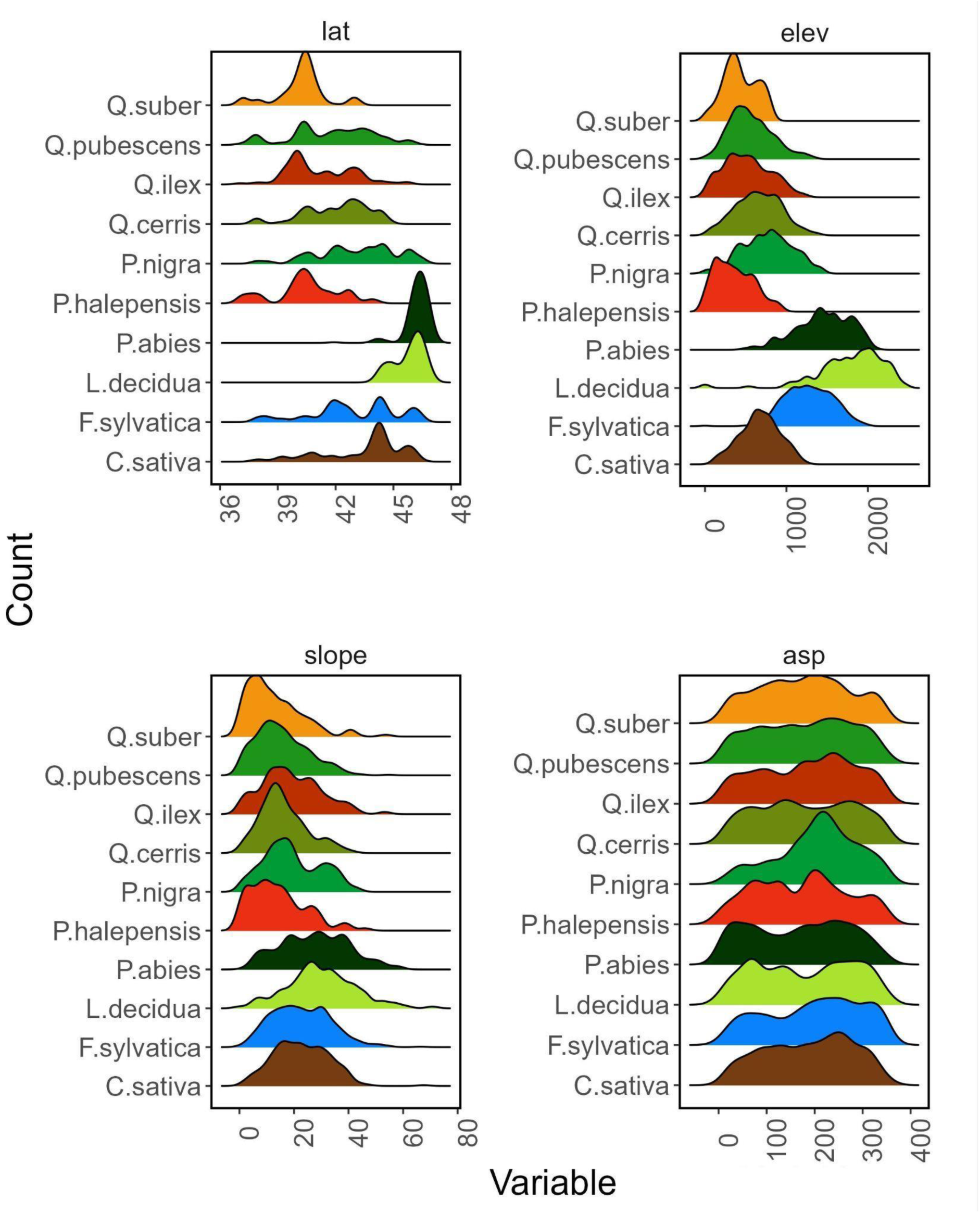
Distribution of auxiliary variable (x-axis) for each selected species (y-axis).

### Statistical analysis

To aggregate VPP over years and ensure we do not miss information from years that may have strongly impacted all species studied, we test for statistical yearly differences in VPP parameters for each species using multiple statistical tests. First, for each year, we test all VPP parameters for normality and homogeneity of variance using the Shapiro-Wilk and Levene’s tests at the 95% confidence level. Second, we fit a one-factor ANOVA with year as the factor to test for overall differences in the VPP distributions. If differences were present, we tested all pairwise comparisons between years using the post hoc general linear hypothesis test (GLHT) at the 95% confidence level. Using the matrix of parametric contrasts for the year factor (calculated via Tukey’s method; Bretz et al., 2010), the GLHT evaluates multiple hypotheses concerning the specified linear function of interest. To assess changes in VPP over the years, we applied the Dunnett post hoc test at a 95% confidence level (Dunnett & Tamhane, 1991), comparing each subsequent year with the first available year in the dataset (i.e., 2017), adjusting the p-value based on the joint normal distribution of the linear function. When Levene’s test detected heteroscedasticity, we used a sandwich estimator (Zeileis, 2006) in both post hoc tests to obtain a heteroscedasticity-consistent covariance matrix.

### Statistical analysis results

Results for the phenological parameters are reported in comparison within each species and across different years. The GLHT results show no statistically significant differences across years for most species and VPP parameters. Figures S2 and S3 present the GLHT post hoc results for the SOSD. C. sativa and F. sylvatica were exceptions, showing statistically significant differences among most year combinations, particularly in amplitude and productivity products (AMPL, SPROD, TPROD, MAXV, and LSLOPE). Interestingly, the Dunnett test revealed fewer differences between 2017 and all other years. Most statistically significant differences were observed across year combinations. Oak species, particularly those from the Mediterranean region (Q. suber, Q. ilex, and Q. pubescens), show the opposite trend, with the largest differences observed between 2017 and other years. After 2017, most VPP parameters remained stable until 2023. Conifer species were the most stable, with Pine species (P. halepensis and P. nigra) showing no statistically significant difference among years in any of the VPP parameters, while L. decidua and P. abies differed in AMPL, SOSD, MAXD, and MINV, particularly between 2017 and 2021 and between 2021 and 2023.

**Figure S2.**
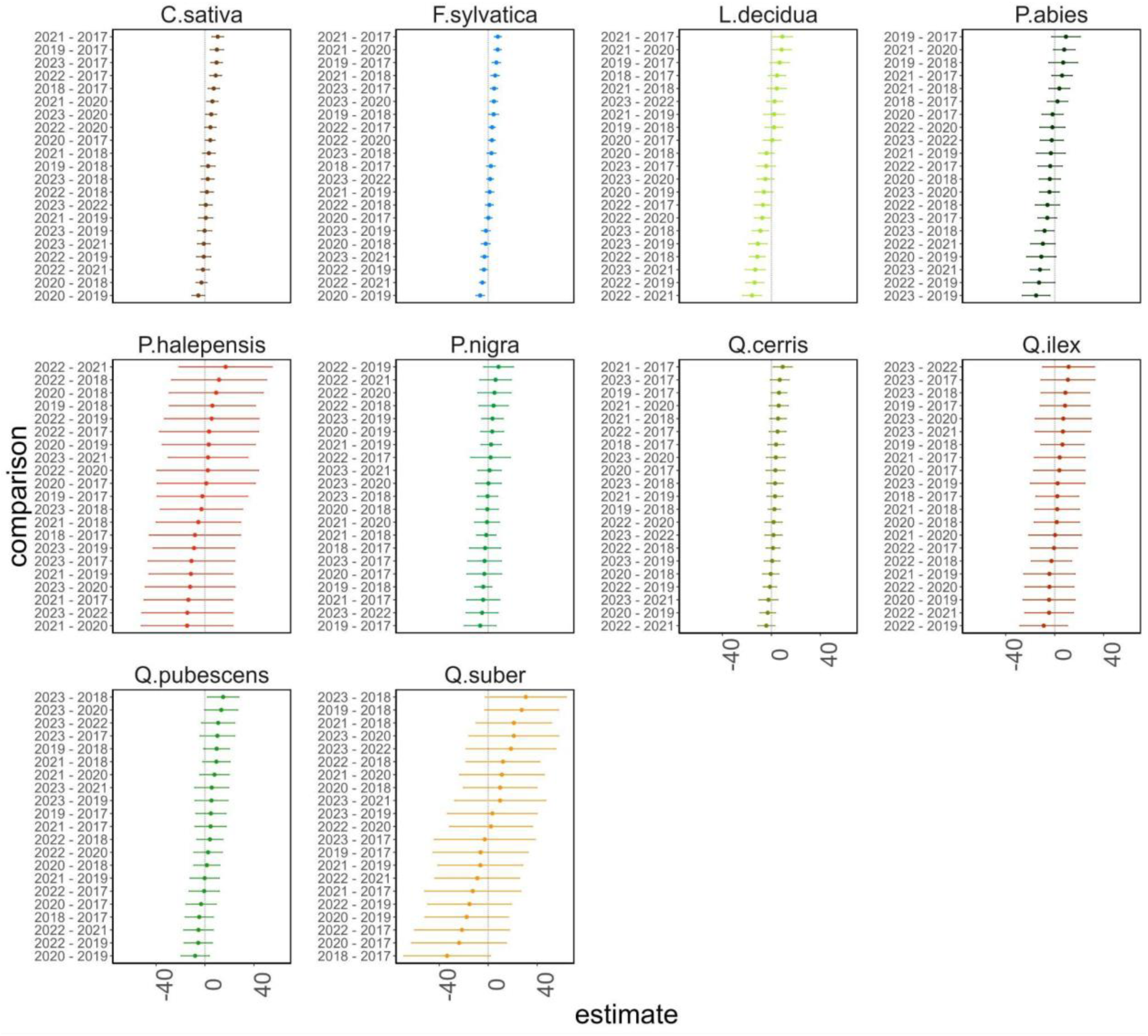
Tukey’s post-hoc test results for each species and all possible pairwise combinations among years for the SOSD variable. Segments that do not cross the vertical line (x = 0) represent statistically significant differences for the year combination in the y-axis.

**Figure S3.**
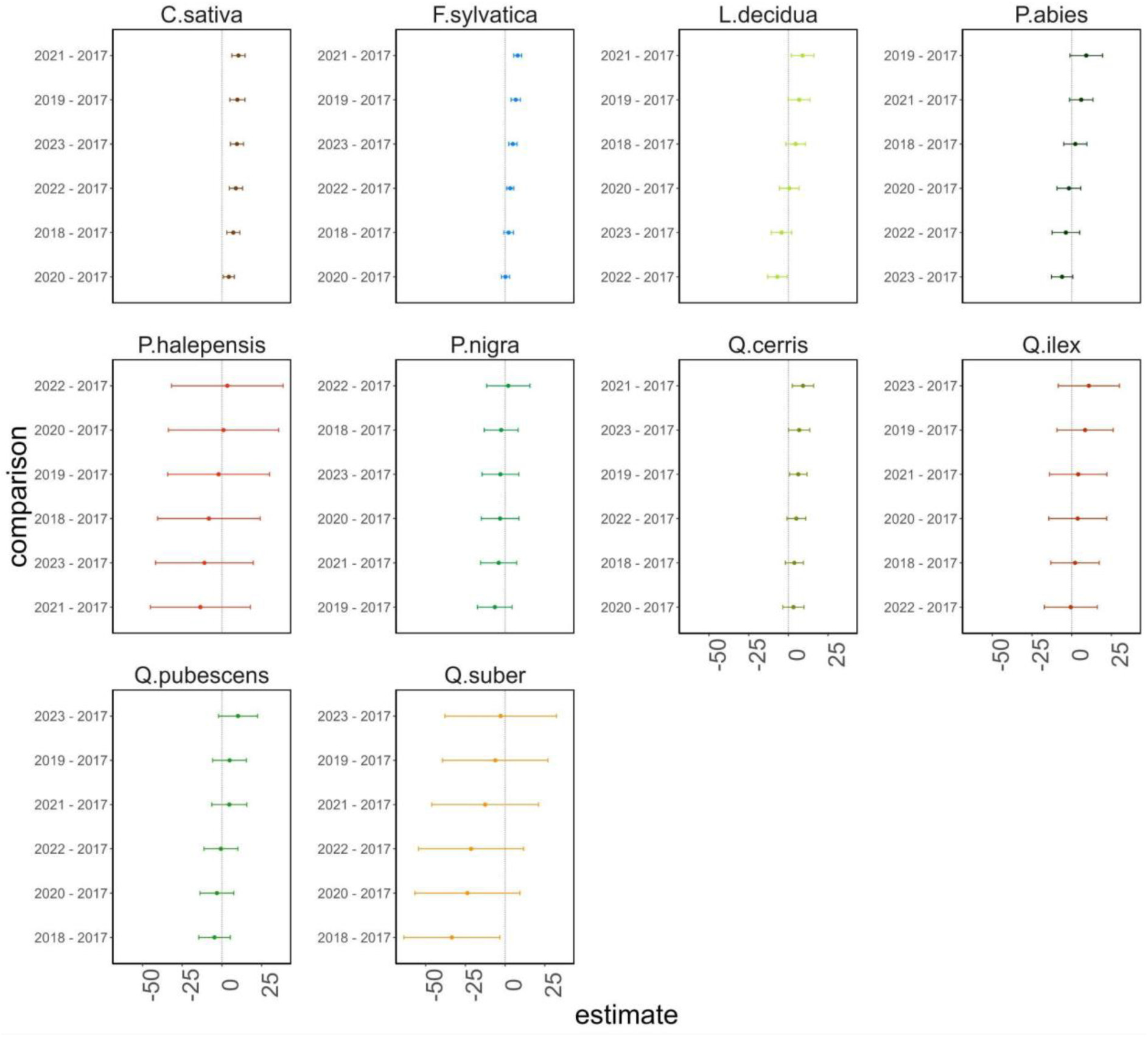
Dunnett’s post-hoc test results for each species and all possible pairwise combinations between 2017 and all other years for the SOSD variable. Segments that do not cross the vertical line (x = 0) represent statistically significant differences for the year combination in the y-axis.

In general, the estimated differences between the means (x-axis in Figs. S2 and S3) and their 95% confidence interval bounds were greater in Mediterranean species than in mountain and Alpine ones. P. halepensis and Q. suber exhibit the highest estimated differences among years with the widest confidence interval. For most species, a positive difference in the SOSD (DoY of the start of the season) compared to the 2017 SOSD coincides with a delayed end of the growing season (higher EOSD), and vice versa. However, this phenomenon is not always followed by a change in the LOS. In Mediterranean species, vegetative season length is essentially fixed, regardless of the degree of SOSD delay. This is not the case for mountain and Alpine species, for which a delayed SOSD shortens their length. However, it is worth noting that the relationships between the start, length, and end of the season and productivity (such as TPROD, SPROD, and MAXV) are complex and not always linear, depending on various factors and interactions. For example, C. sativa exhibits a statistically significant delay in SOSD from 2018 to 2023 compared to 2017, but no difference in TPROD. In contrast, Q. suber exhibits the opposite pattern, with no difference in SOSD relative to 2017 but significant positive differences in TPROD.

**Figure S4.**
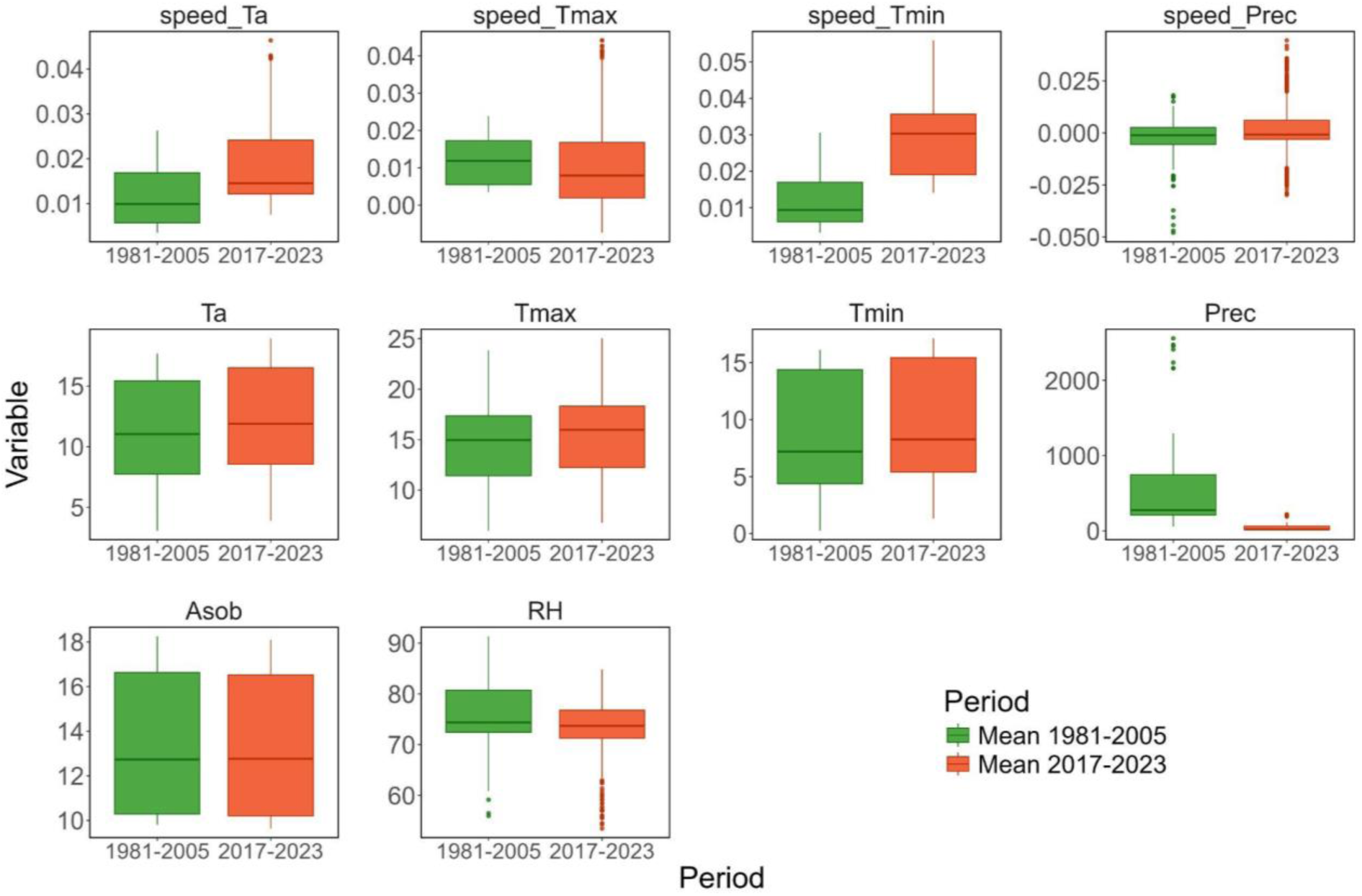
Annual climate metrics distribution comparison: green boxplots represent the climatic variables’ mean of all field plots over the period 1981-2005, while the red ones represent the mean over the period 2017-2023 (covered by this study). Speed represents the yearly rate of increase for that variable. Variables are: mean, maximum and minimum temperature (Ta, Tmax, Tmin), precipitation (Prec), solar radiation (Asob), and relative humidity (RH).

